# Sexual and developmental variations of ecto-parasitism in damselflies

**DOI:** 10.1101/2021.12.06.471459

**Authors:** Shatabdi Paul, Md Kawsar Khan, Marie E. Herberstein

## Abstract

The prevalence and intensity of parasitism can have different fitness costs between sexes, and across species and developmental stages. This variation could arise because of species specific sexual and developmental differences in body condition, immunity, and resistance. Theory predicts that the prevalence of parasitism will be greater in individuals with poor body condition and the intensity of parasitism will be greater in individuals with larger body size. These predictions have been tested and verified in vertebrates. In insects, however, contradictory evidence has been found in different taxa. Here, we tested these predictions on two species of *Agriocnemis* (*Agriocnemis femina* and *Agriocnemis pygmaea*) damselflies, which are parasitized by *Arrenurus* water mite ectoparasites. We measured body weight, total body length, abdomen area and thorax area of non-parasitized damselflies and found body condition varied between males and females, between immature females and mature females and between *A. femina* and *A. pygmaea*. Then, we calculated the parasite prevalence, i.e., the frequency of parasitism and intensity, i.e., the number of parasites per infected damselfly in eleven natural populations of both species. In line to our predictions, we observed greater prevalence in immature females than mature females but found no difference in parasite prevalence between males and females. Furthermore, we found that parasite load was higher in females than males and in immature females than mature females. Our result also showed that the frequency and intensity of parasitism varied between the two studied species, being higher in *A. pygmaea* than *A. femina*. Our study provides evidence that parasitism impacts sexes, developmental stages and species differentially and suggests that variation may occur due to sex, developmental stage, and species-specific resistance and tolerance mechanism.

## Introduction

According to sexual selection and resource allocation theory, individual differences in allocating resources to reproduction, immunity, and survivability may explain the variation of parasitism between sexes [1–3]. Males under strong sexual selection may invest more resources in mating related traits such as conspicuous color, elaborated mating display, and less towards immunity which increases their susceptibility to parasite infection [1,2,4]. On the other hand, fecundity driven selection on females may result in greater allocation of energy to fecundity related traits thereby maintaining larger body size, and higher fat content [5–9] and less to immunity and resistance. This could increase the chance of being infected, and can provide more nutrition reservoir and larger surface area for a greater number of parasites in females [10–14]. Furthermore, sex specific behaviour, such as habitat use, time spend in search of mates, sex specific life history traits such as developmental rate, and variation in stress levels imposed by mating system may also contributes to the variation in parasitism between the sexes [5,13–18].

Physiology, immunity and behavior can also vary across populations, developmental stages, and among different species which may contribute to inter- and intraspecific variation of parasitism as well as variation between immature and mature individuals [9,19–23]. For example, intraspecific variations of parasitism were studied in *Dineutus nigrior* beetles, *Paratrichocladius rufiventris* midges, Clinocerinae flies, and *Ranatra chinensis* water scorpions where larger individuals, and individuals with lower immunity were shown to be more susceptible to parasites [7,13,24–27]. On the other hand, juvenile individuals typically have a smaller body size and may have an underdeveloped immune system, which might contribute to a greater susceptibility of parasitism during immature stages. For instance, in *Appasus japonicus* bugs, juveniles showed higher parasite loads than adults [26]. However, parasite prevalence was not different between developmental stages [26].

Odonates (dragonflies and damselflies) are frequently parasitized by protozoan endoparasites and *Arrenurus* mite ectoparasites [28,29]. The frequency and intensity of parasitism has been shown to vary among species, across populations, developmental stages, and between sexes [3,9,26,30]. Individual differences in physiology (immune resistance and tolerance), habitat use and behavior accounted for the interspecific variation of parasitism in *Nehalennia irene, Calopteryx maculata, Enallagma chromatallagma, Ischnura posita* and *Lestes forcipatus* damselflies [31–36]. The impact of host sex on the likelihood of being parasitized is contested. Some studies found higher parasite infections in males than female [3,12,37–39]; whereas others found the opposite with a female bias in parasite prevalence and intensity [38,20,40–43]. Whereas, no sex differences in parasite prevalence and intensities were found in *C. eponina* and *P. longipennis* damselflies [44– 47]. The conflicting evidence in different species suggests that variation of parasitism between sexes maybe species specific.

There is evidence that the frequency and intensity of parasite infections varies among different developmental stages of damselflies [48]. Hecker *et al*. (2002) reported that larval damselflies had a higher gregarine prevalence and intensity than newly emerged damselflies [20]. Conversely, final instar damselfly larva of *Lestes forcipatus* damselfly carried more water mites than late instar larvae [12]. Rates of parasitism are expected to vary between sexually immature and mature stages of damselflies [9,20,22]. Nevertheless, to the best of our knowledge, no studies have explored the variation of parasitism between sexually immature and mature damselflies.

Here we aim to determine variation in parasite prevalence and intensity between sexes, and developmental stages in *Agriocnemis femina* and *A. pygmaea* damselflies. The rate and extent of parasitism is likely to depend on the sex specific condition of damselflies with a larger surface area and higher nutrition providing more space and resources for parasites. To test this idea, we conducted morphometric measurements including body weight, total body length, abdomen area and thorax area of non-parasitized damselflies to determine whether body condition varies with sex, developmental stages and species. We found female damselflies were heavier and had greater surface area compared to males. Moreover, immature females are smaller and lighter than mature females. Finally, *A. femina* were larger and heavier compared to *A. pygmaea*. Based on these morphometrics, we predicted that (1) parasite numbers would be higher in female damselflies than males, because females are larger and had greater surface area, (2) sexually mature females would have greater parasite loads because of their larger surface area and greater resource indicated by heavier body weight and (3) *A. femina* would be parasitized more frequently and with a larger number of parasites than *A. pygmaea* because of their larger body size and body weight. Our data also address alternative hypotheses that might support the sex and developmental dependent immune performance. Accordingly, male damselflies would be more parasitized than females, because females tend to have better immunity than males [11], immature females would have greater parasite prevalence because of their underdeveloped immune system compared to adults. We calculated frequency of parasitism and intensity of parasitism across 11 natural populations to test our predictions.

## Materials and methods

### Study systems

*Agriocnemis femina* and *Agriocnemis pygmaea* are small damselflies (wing size 10.5-11.00 mm, and wing size 9.75-11.5 mm respectively) of the Coenagrionidae family [49,50;Figs 1a and 1b]. These species are widely distributed in South Asia, South East Asia, and Australia [50–51]. They are commonly found in grasslands associated with ponds, lakes, marshes and in paddy fields, and cohabit with *Ceriagrion coromandelianum, A. kalinga A. lacteola, and Orhterum sabina* [49]. The males of *A. femina* are differentiated from other sympatric species by the size and shape of anal appendages (Epiproct larger than cerci) [50;Fig 1c]. *Agriocnemis pygmaea* males are distinguished by their smaller size, green ante-humeral stripes, orange abdomen tip, and larger cerci than epiproct [50;Fig 1f]. *Agriocnemis femina* females are recognized by their protruded ridge on the prothorax [50;Figs 1a and 1b] whereas *A. pygmaea* females are identified by the pink ante-humeral stripe in juveniles and green ante-humeral stripes in adults (Figs 1d and 1e). Females of both species exhibit ontogenetic colour change from red to green which signals sexual maturity and reduce pre-reproductive male mating harassment [49]. Males on the other hand, do not undergo such conspicuous changes therefore, sexually immature and mature males cannot be identified accurately in the field.

**Fig 1.**
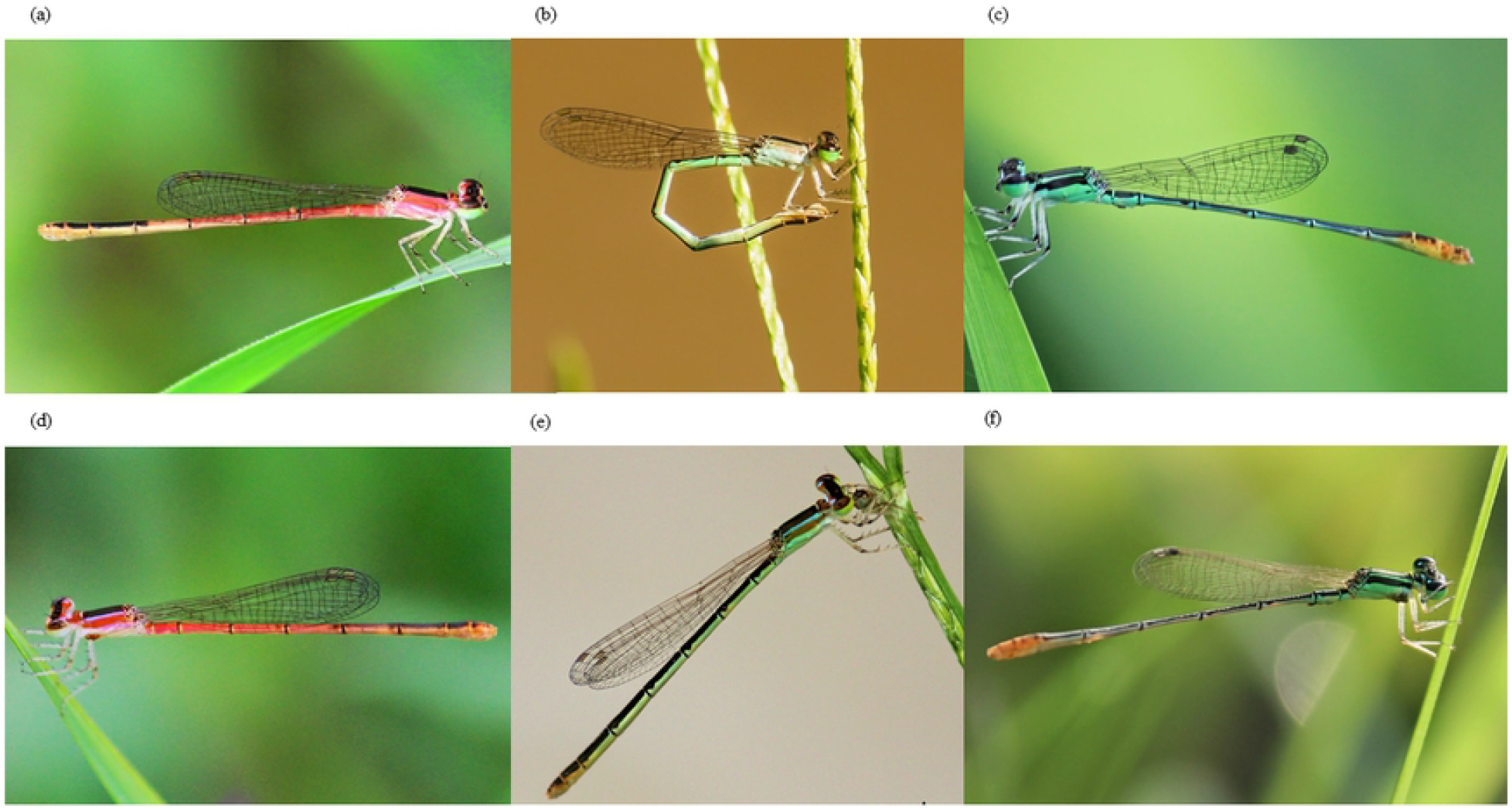
Photographs of females, males, immature and mature females of *Agriocnemis femina* and *Agriocnemis pygmaea* respectively. (a) Photograph of an immature female *Agriocnemis femina*. (b) Photograph of a mature female *Agriocnemis femina*. (c) Photograph of a male *Agriocnemis femina*. (d) Photograph of an immature female *Agriocnemis pygmaea*. (e) Photograph of a mature female *Agriocnemis pygmaea*. (f) Photograph of a male *Agriocnemis pygmaea*.

### Study sites

*Agriocnemis femina* is a common damselfly in the northeastern region of Bangladesh [53,54], while *A. pygmaea* is commonly found in central and southwestern region of Bangladesh [49,55]. *Agriocnemis femina* is seen in flight throughout the year, although the number of individuals peaks between April to July [50]. The flight period of *A. pygmaea* is all year round, but the population level peaks between June to August [56]. We conducted field studies for *A. femina* in six different sites in the northeastern region of Bangladesh between March 2021 to June 2021 (Table 1). *Agriocnemis pygmaea* were studied in five field sites in the central and southwestern region of Bangladesh between May 2017 to June 2017 (Table 1). No permission was required to collect specimens because none of the two species are endangered in Bangladesh and field sites were not located in protected areas.

**Table 1.**
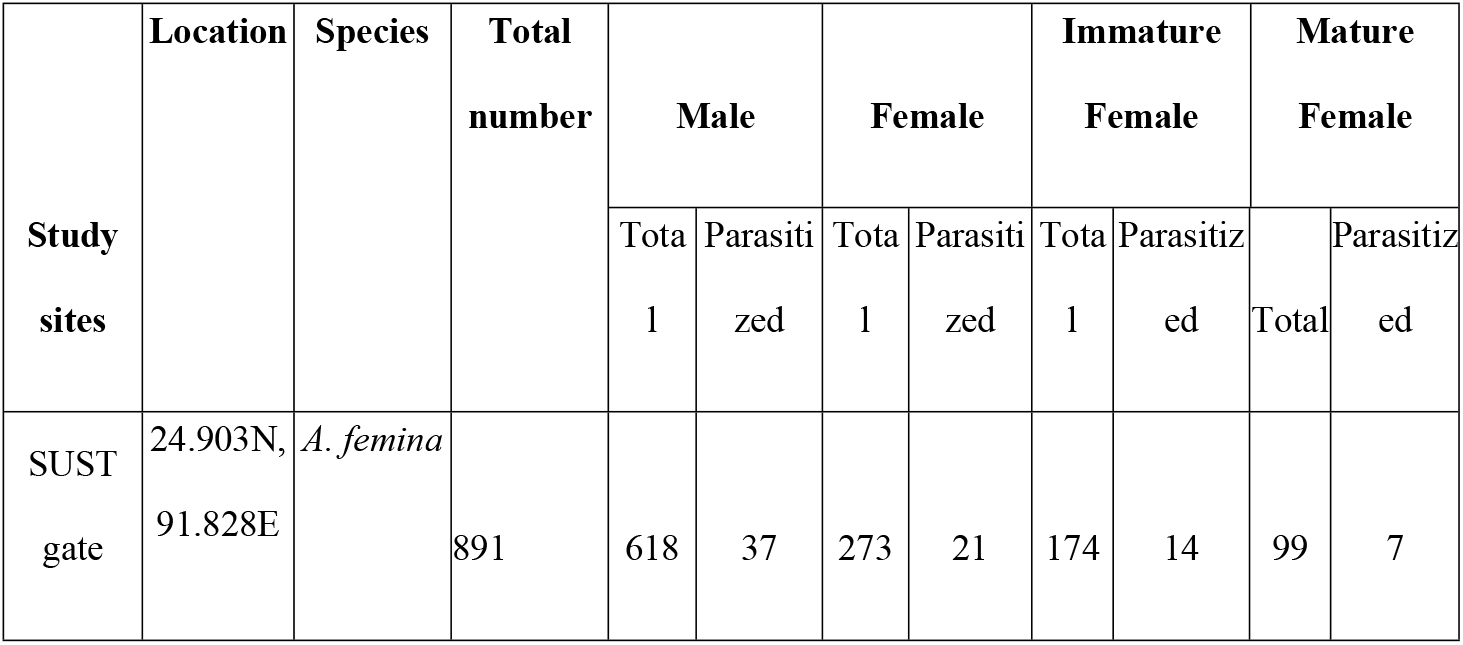
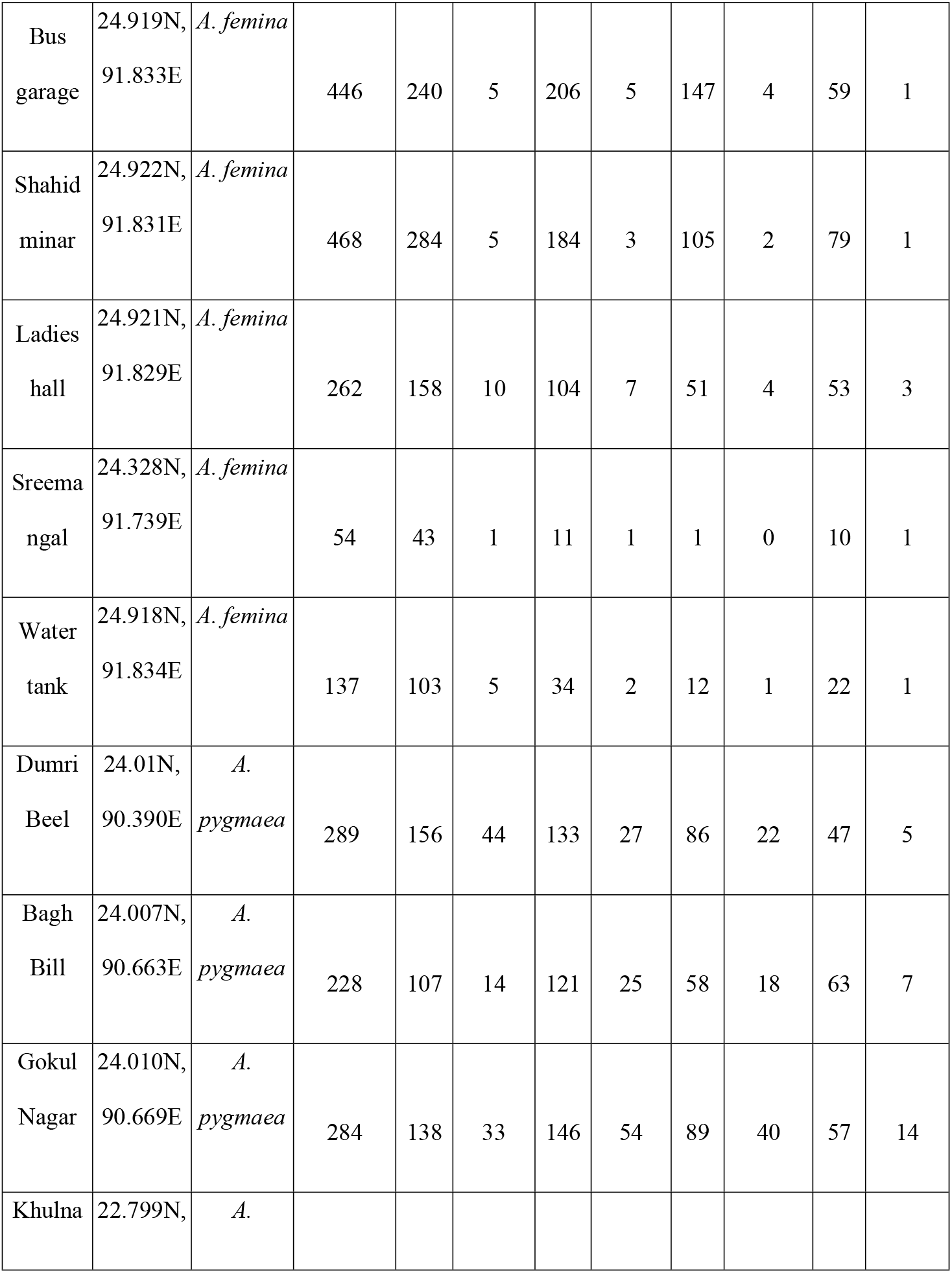
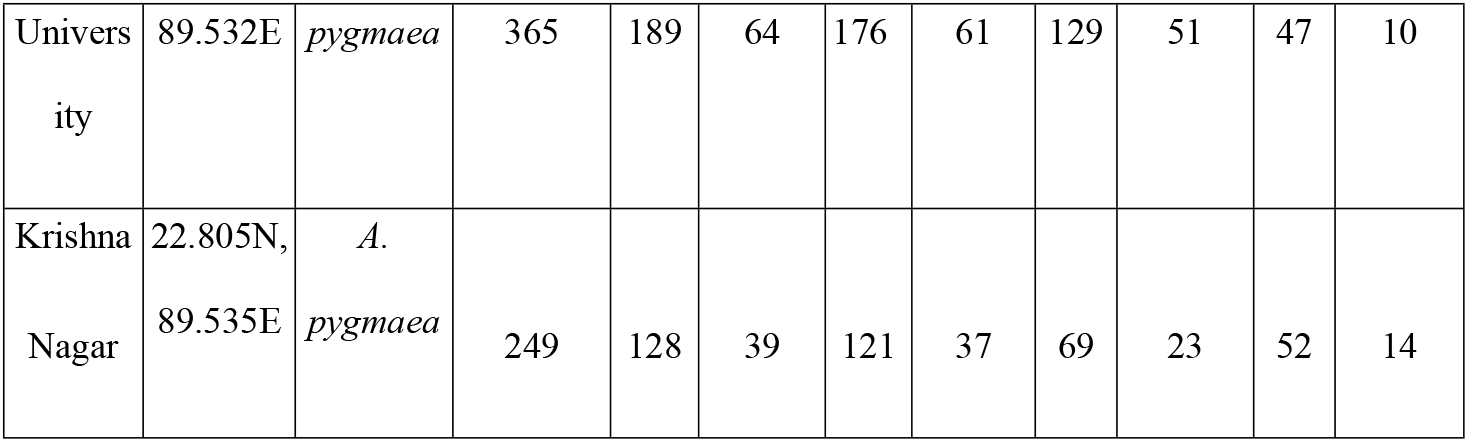
Study sites, locations and number of total and parasitized *Agriocnemis femina* and *Agriocnemis pygmaea* captured to calculate parasite prevalence.

### Ectoparasite mite

Water mites of Arrenuridae family are the most common ectoparasites of damselflies [29,57,58]. The water mites infect damselflies when the adult emerges from the larval stage or in a later life stage when the adults visits waterbodies for foraging or ovipositing [58,59]. The ectoparasite mites attach themselves to the exocuticle of the host damselflies and form a feeding tube to extract body fluids of the host [60–62]. Water mites depend on host damselfly nutritional resources for their growth and development [63].

### Morphometric analysis

We collected damselflies from the field using an insect sweep net and stored them in 95% ethanol. We transported the damselflies back to the laboratory for morphometrics analysis. We then measured body weight, total body length, abdomen area and thorax area of damselflies. First, we placed the damselflies on an absorbent paper for exactly two minutes to evaporate the ethanol [49] and then weighed the damselfly on a balance (Shimadzu ATY 224 electronic balance, Shimadzu Corporation, Japan). Next, we positioned the damselflies laterally along with a scale and took photographs. We measured total body length, and abdomen length, width of the fifth abdominal segment, length and width of thorax of the damselflies from the digital photographs using the ImageJ software [64]. Later, we calculated abdomen area and thorax area using equation (area = 1/2 × width × π × length) [49].

### Frequency and intensity of parasitism

We captured damselflies with an insect sweep net while walking along the edge of ponds and submerged grasslands by following previously established method [65]. We recorded the sex (male or female) for every captured individual and developmental stage of the females (immature or mature). We visually inspected the captured specimens for a presence of parasites. For parasitized individuals, we counted the number of parasites. After inspection, we marked the damselflies on their wings with a permanent marker and released them into their natural population. This marking procedure avoids the recapture of the same individual [66,67]. We calculated the frequency of parasitism, i.e., the proportion of individuals parasitized, and the intensity of parasitism, i.e., the number of parasites present for each parasitized individual. We conducted the fieldwork between 08:00 and 10:00 hours when the species are mostly active, mating occurs, and condition are favorable for field work.

### Statistical Analyses

We applied linear mixed effects models (LMMs) to determine whether body weight, total length, abdomen area and thorax area varies between non-parasitized males and females, immature and mature female damselflies. We fitted LMMs with body condition using sex and developmental stage as fixed factors and species as random factor. We used r.squaredGLMM function of the R package “MuMIn” to determine the effect size of the models [68]. We further applied Mann-Whitney *U* test to determine whether body weight, total length, abdomen area and thorax area varies between *A. femina* and *A. pygmaea*.

We used generalized linear mixed models (GLMMs) with binomial distributions to determine whether males are more frequently parasitized than females. We fitted GLMMs with parasite frequency as a response variable, sex and species as fixed effects and study site as a random factor (model: glmer (cbind (parasitized, total-parasitized) ∼ sex + species + (1|study_site), family = binomial)). To determine whether immature females are parasitized more frequently than mature females, we fitted GLMM with parasite frequency as a response variable, developmental stage as a fixed effect, and study site as a random effect (model: glmer (cbind (parasitized, total-parasitized) ∼ developmental stage + (1|study_site), family = binomial)).

We applied generalized linear mixed models (GLMMs) with a poisson distribution to test whether parasite intensity differs between sexes, and the two study species. We fitted GLMM with parasite intensity as a response variable, sex and species as fixed factors, and study site as a random effect (model: glmer (parasite number ∼ sex + species + (1|study_site), family = poisson)). We further applied GLMM to determine whether parasite intensity varies between immature and mature female developmental stages in parasite intensity. The full model was glmer (parasite number ∼ developmental stage + (1|study_site), family = poisson). We used the r.squaredGLMM function of the “MuMIn” R package to determine the effect size of the models [69]. We analyzed all the data in the R version 4.0.3 using the ‘lme 4’ [69] and ‘MuMIn’ [70] packages. All values are estimate ± standard error.

## Results

### Morphometric analysis

Females and males differ significantly in their body weight, total length, and body surface area. Females had significantly higher body weight, total body length, abdomen area and thorax area than males (all *P* < 0.0001; Fig 2, S1 Table). Body weight, total length and body surface area were also different between the developmental stages. Immature females had lower body weight, body size, abdomen area, thorax area compared to mature females (all *P* < 0.1; Fig 2; S2 Table). *Agriocnemis femina* had greater body weight, body size and thorax area than *A. pygmaea* (all P < 0.0005; Fig 2; S3 Table). However, *A. femina* and *A. pygmaea* did not significantly vary in abdomen area (*P* = 0.1; Fig 2; S3 Table).

**Fig 2.**
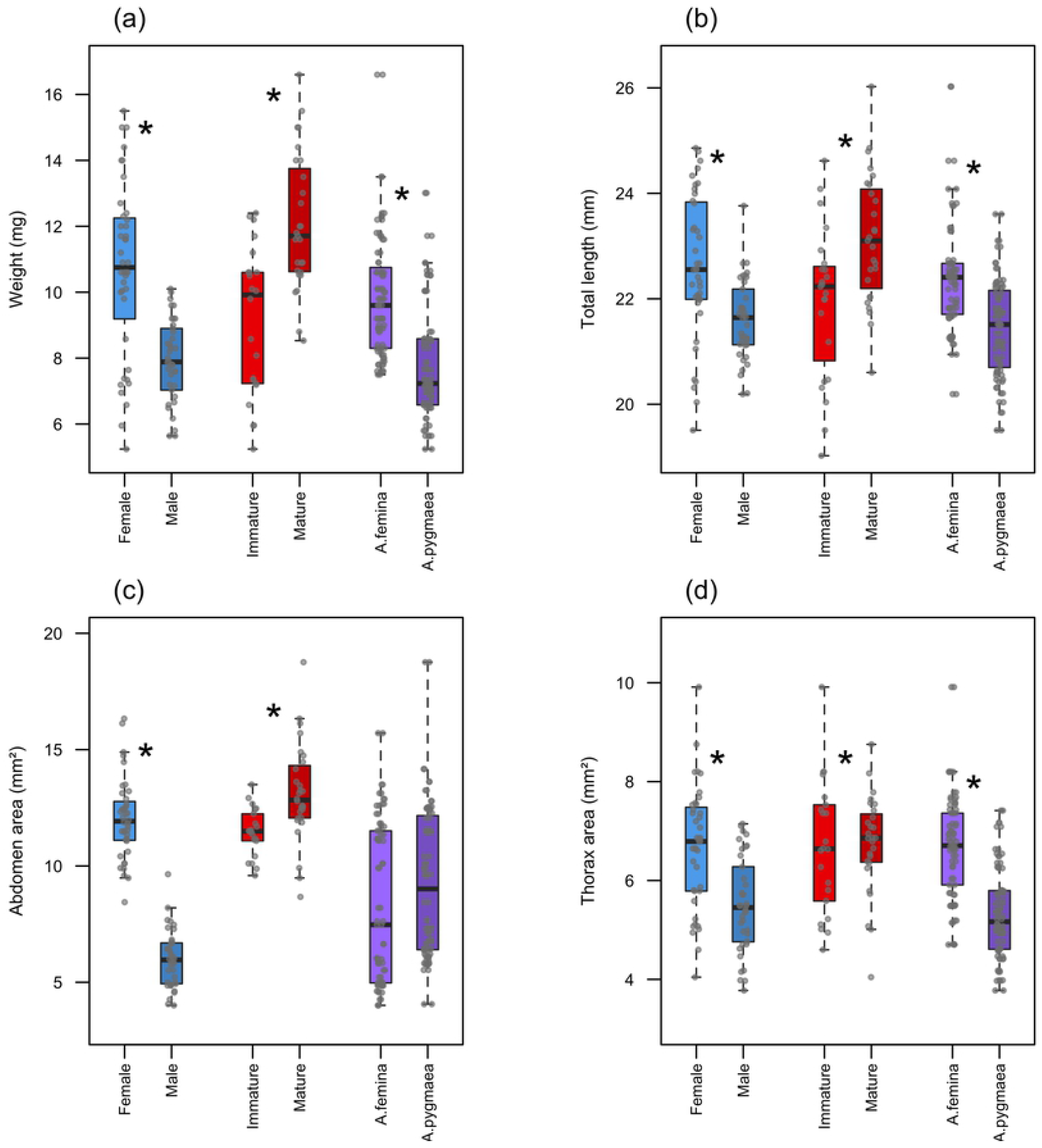
Morphometric measurements of non-parasitized and fresh females, males, immature and mature females of *Agriocnemis femina* and *Agriocnemis pygmaea* respectively. (a) Body weight of females and males; immature and mature females; *Agriocnemis femina* and *Agriocnemis pygmaea*. (b) Total length of females and males; immature and mature females; *z femina* and *A. pygmaea*. (c) Abdomen area of females and males; immature and mature females; *A. femina* and *A. pygmaea*. (d) Thorax area of females and males; immature and mature females; *A. femina* and *A. pygmaea*. The boxplots indicate the median, 25th and 75th percentiles. The error bars extend downward from the first quartile to the minimum and upward from the third quartile to the maximum data points. Data points which are 1.5 times greater than the interquartile range are excluded. * symbolizes significant variation between studied groups.

### Frequency and intensity of parasitism

A total of 3673 individuals were inspected (N = 2258 *A. femina* and N = 1415 *A. pygmaea*) of which 16.33% were parasitized (4.52% of *A. femina* and 28.14% of *A. pygmaea* (Table 1)). Parasite prevalence did not differ significantly between males (N = 2164) and females (N = 1509) (GLMM: estimate = -0.1373 ± 0.1039, *Z* = -1.322, *P* = 0.186; *R*^*2*^ = 0.986; Fig 3a). Parasite prevalence was significantly higher in immature females (N = 921) than in mature females (N = 588) (GLMM: estimate = -0.7328± 0.1669, *Z* = -4.391, *P* < 0.0001; *R*^*2*^ = 0.967; Fig 3b). Furthermore, parasite prevalence was significantly higher in *A. pygmaea* compared to *A. femina* (GLMM: estimate = 2.1815 ± 0.27, *Z* = 7.924, *P* < 0.0001; *R*^*2*^ = 0.986; Fig 3c).

**Fig 3.**
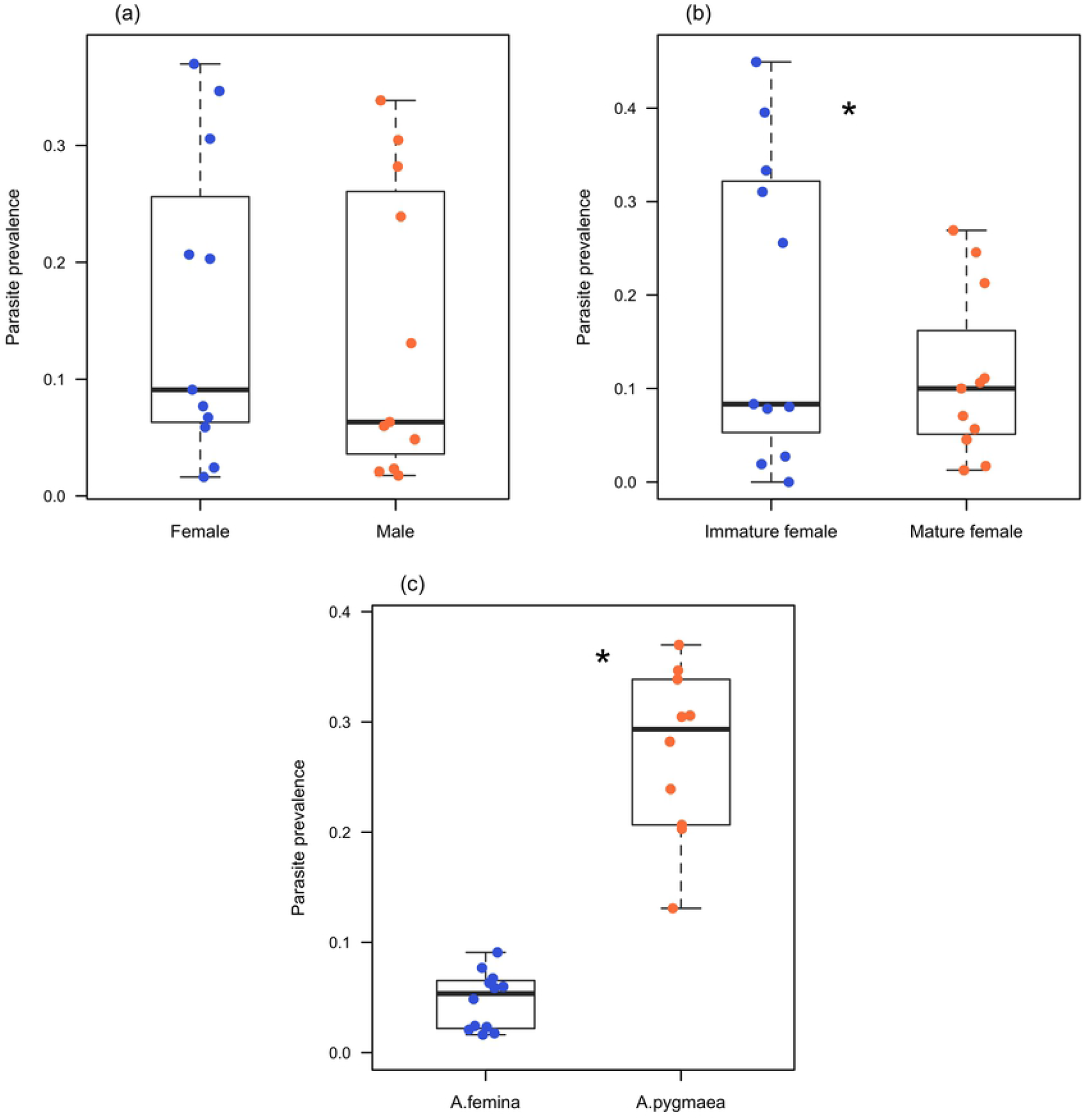
Parasite prevalence in females, males, immature and mature females of *Agriocnemis femina* and *Agriocnemis pygmaea* respectively. (a) Parasite prevalence in females and males. (b) Parasite prevalence in immature and mature females (c) Parasite prevalence in *Agriocnemis femina* and *Agriocnemis pygmaea*. The bold lines within boxes depict the median. The bottom and top borders of the boxes indicate 25th and 75th percentiles, respectively. The whiskers include all data points excluding the data that are beyond 1.5 times the interquartile range. Circle indicates parasite prevalence at each study site. * denotes p <0.0001.

Parasite number varies from 1 to 14 in infected individuals with a mean of 3.429 ± 0.0959. Parasite intensity was significantly lower in male damselflies than females (GLM: estimate = -0.18542 ± 0.04323, *Z* = -4.289, *P* < 0.0001; *R*^*2*^ = 0.139; Fig 4a). Moreover, parasite intensity was significantly lower in mature females than immature females (GLM: estimate = -0.32078 ± 0.07787, *Z* = -4.119, *P* < 0.0001; *R*^*2*^ = 0.1967; Fig 4b). Parasite intensity was greater in. *A. pygmaea* than *A. femina* (GLM: estimate = 0.58565 ± 0.09055, *Z* = 6.468, *P* < 0.0001; *R*^*2*^ = 0.139; Fig 4c).

**Fig 4.**
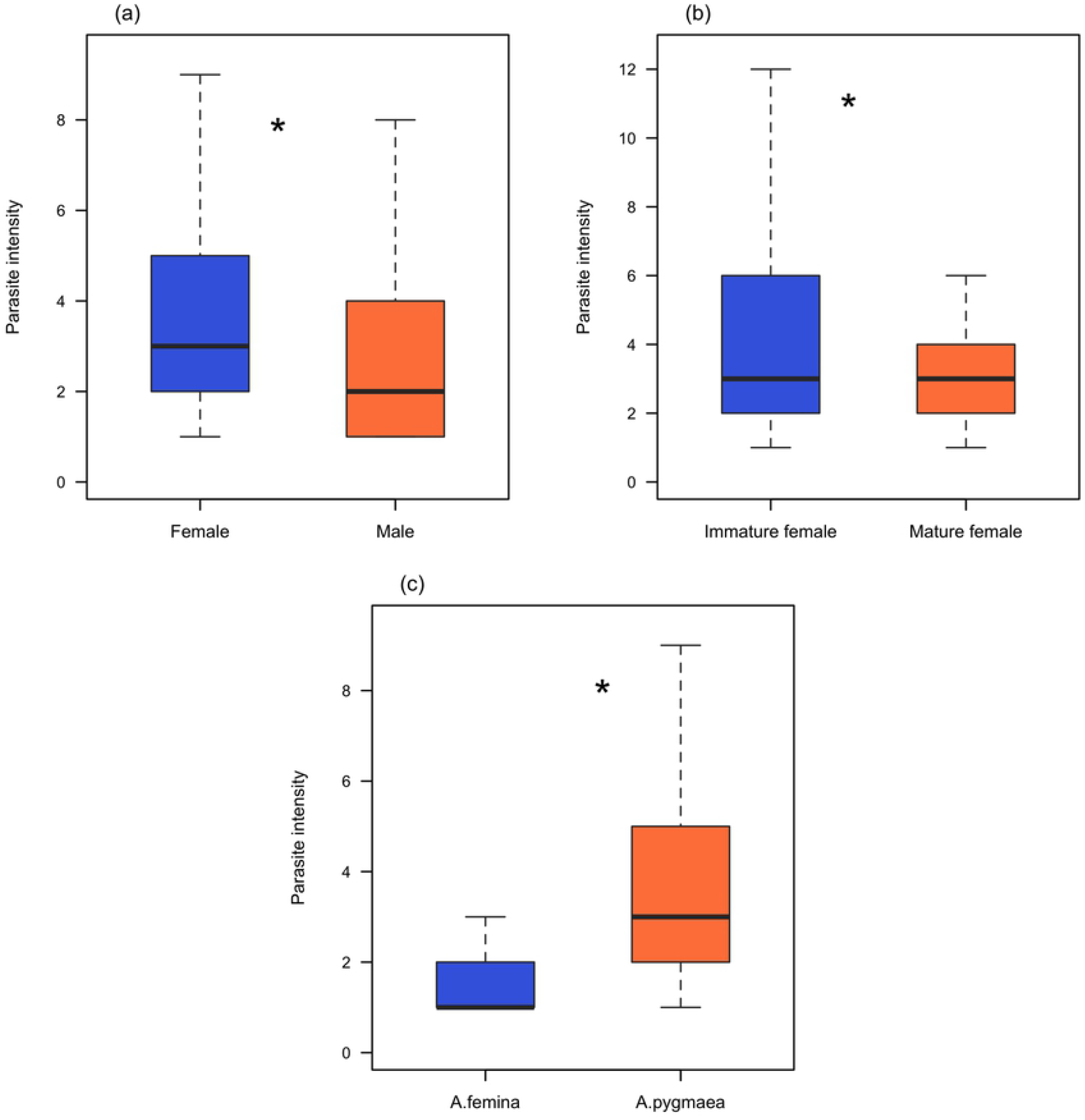
Parasite intensity in females, males, immature and mature females of *Agriocnemis femina* and *Agriocnemis pygmaea* respectively. (a) Parasite intensity in females and males. (b) Parasite intensity in immature and mature females. (c) Parasite intensity in *Agriocnemis femina* and *Agriocnemis pygmaea*. The boxplots represent the median, 25th and 75th percentiles. The error bars extend downward from the first quartile to the minimum and upward from the third quartile to the maximum data points. Data points beyond the error bars are 1.5 times greater than the interquartile range and excluded. * indicates significant variation (p <0.001) between studied groups.

## Discussion

Parasite frequency and intensity may vary between males and females, and developmental stages of a species because of differences in morphology, physiology, behavior, and immunity. Here, we determined sexual and developmental variation of morphological traits and intensity and frequency of parasitism in two damselfly species. We found that females were larger, heavier and had greater surface area than males. Mature females also had a larger body size, more body weight, and greater surface area than immature females. Our results showed that, parasite prevalence did not vary between sexes, but females had significantly higher parasite intensity compared to males. We further showed that parasite prevalence and parasite intensity were greater in immature females than mature females. Finally, we also found interspecific variation of parasitism between the two studied species, *A. pygmaea* had higher frequency and intensity of parasitism than *A. femina*.

Contrary to our prediction, we did not find a significant difference in parasitism between males and females. Behavioral and life history trait differences can contribute to distinct risks of parasitism for males and females. For example, in species where eggs are oviposited directly into the water or on vegetation on or in the water, females spend more time in or near water than males [71]. During ovipositing, females suffer increased exposure to ectoparasites and consequently increased risk of parasitism. Accordingly, female biased parasite prevalence was observed in *Lestes sponsa, Coenagrion pulchellum*, and *Ischnura verticalis* were ovipositing requires contact with water [8,29,43]. By contrast, *A. pygmaea* and *A. femina* females oviposit on grasses 20-30 cm above the water line (MKK personal observation; Figs 1b and 1e), which reduces contact with water during oviposition and might subsequently result in lower parasite exposure. This might explain the lack of female biased parasitism. Furthermore, immunity and resistance against parasites might not vary between males and females [29]. Further studies are required to understand the impact of sex specific immunity on parasitism in *A. pygmaea* and *A. femina*.

The lack of any sexual bias in the prevalence of parasitism in our study species concords with previous studies in damselflies i.e., *Lestes disjunctus, I. verticalis, Nehalennia Irene*, and other arthropod hosts (e.g. crustaceans, coleopterans, dipterans) that did not find differences in parasitism between males and females [72–79]. These results suggest that unlike vertebrates, sex differences in parasitism might not be the norm in invertebrates [5].

We showed that parasite intensity was greater in females than males. Female biased sexual size dimorphism and better condition in females in *A. pygmaea* and *A. femina* most likely contributed to the increased parasite intensity in females [49]. The higher body mass of females could provide greater resources to parasites therefore could harbor more parasites [80]. Larger body size can also provide greater surface area for parasite accumulation [74]. The larger body size in *Agriocnemis* females and greater thoracic and abdominal area therefore probably accounts for the greater parasite intensity. Furthermore, due to selection on fecundity, females may invest more resources to the quality and quantity of eggs, and less in immunity [1,3,10,81]. Thus, higher parasite load in females could be the result of a trade-off between fecundity and immunity investment [9]. In accordance to our findings, higher parasite intensity in females have been shown in other damselflies *Ischnura elegans, Coenagrion pulchellum, Lestes disjunctus*, and in *Dineutus nigrior* whirligig beetle [9,13,29,60,82].

Physiology and behavior of damselflies vary between different development stages, consequently, immature and mature individuals may differ in parasite prevalence and intensity [83,84]. We found significantly higher prevalence and intensity in immature females compared to mature females. While immature female damselflies are less active than mature females [85,86], they spend longer periods of time at water-side vegetation after emergence as shown in *Enallagma boreale, E. ebrium, E. aspersum, Ischnura elegans, Xanthocnemis zealandica*, and in *Lestes disjunctus* [19,20,48,82,87,88]. Whether these behavioural differences contribute to higher parasite frequency in immature *Agriocnemis* is unclear.

Furthermore, immunological variation can also contribute to the observed variation of parasitism between developmental stages. For example, less pronounced immunity in immature individuals of *Enallagma ebrium, Coenagrion hastulatum*, and *Ischnura elegans* have been found to increase parasitism [9,48,89].Whether behavioral, nutritional, or immunological variation, or a combination of these contribute to the variation of parasitism between immature and mature females in *Agriocnemis* damselflies is an exciting question of further studies.

Our results showed that, in parasitized individuals, the intensity was higher in mature females than immature females. Females may allocate resources to fecundity traits such fat storage during ontogenesis. *Agriocnemis* females, like other damselflies, acquire body mass, increase body size, and accumulate more fat and protein throughout their development [49,90]. The larger body size, greater surface area, and higher nutritional resources in mature females most likely contribute to the higher parasite load in mature female compared with immature females. Because higher parasite load in individuals is likely to result in greater mortality, immature damselflies parasitized with high parasite load may succumb to death during development. As a result, when we sampled the immature damselflies, those with fewer parasites may have been over-represented.

Interspecific difference in physiology such as body size, and weight can contribute to the species-specific variation of parasitism where larger species are more likely to be heavily parasitized. In contrast, we found higher parasite prevalence and intensity in the smaller and less heavy *A. pygmaea* compared to *A. femina*. In support of this, larger and heavier individuals within a species, have better immunity than smaller and lighter individuals [91,92]. Accordingly, larger species might be able to invest more into immunity compared to smaller species, which could explain why

*A. femina* had lower parasite frequency. Alternatively, the variation of microhabitat and environmental factors might contribute to the variation of parasitism between the two study species. The study species were not sympatric; *A. pygmaea* were sampled from the central and southern region whereas *A. femina* were studied in the northern region of Bangladesh. It is possible that the southern and central region provides favorable condition for the ectoparasites thereby causing higher parasitism in *A. pygmaea*.

## Conclusion

Our study provides new insights into the variation of parasite infection in *Agriocnemis* damselflies indicating that sex and developmental stage bias parasitism. It also offers a comparison of variations in infection among different species of damselflies. This study provides the basis for future work to identify the causes of variation and understand damselflies’ defense mechanism against parasitism.

## Statement of diversity and inclusion

We strongly support equity, diversity and inclusion in science. The authors come from different countries (Bangladesh, Austria and Australia) and represent different career stages (Masters student, Early career researcher, & Professor). One or more of the authors self-identifies as a member of the LGBTQI+ community. One or more authors underrepresented ethnic minority in science. While citing references scientifically relevant for this work, we actively worked to promote gender balance in our reference list.

## Acknowledgment

We acknowledge the *Wallumattagal clan of the Dharug nation* as the traditional custodians of the Macquarie University land. We thank Prof. Dr. Swapan Kumar Sarker for proving logistical support, Mostakim Rayhan for helping with the field work, Department of Biochemistry and Molecular Biology, Shahjalal University of Science and Technology for providing lab access. Authors thanks their families for their support.

## Supporting Information

**S1 Table. Linear mixed effects models (LMMs) for differences in body weight, total length, abdomen area and thorax area between non-parasitized males and female damselflies**. Results from linear mixed effects models (LMMs) for differences in body weight, total length, abdomen area and thorax area between non-parasitized males and female damselflies.

(DOCX)

**S2 Table. Linear mixed effects models (LMMs) for differences in body weight, total length, abdomen area and thorax area between non-parasitized immature and mature female damselflies**. Results from linear mixed effects models (LMMs) for differences in body weight, total length, abdomen area and thorax area between non-parasitized immature and mature female damselflies.

(DOCX)

**S3 Table. Mann-Whitney U-test for differences in body weight, total length, abdomen area and thorax area between non-parasitized *Agriocnemis femina* and *Agriocnemis pygmaea***. Results of Mann-Whitney U-test for differences in body weight, total length, abdomen area and thorax area between *Agriocnemis femina* and *Agriocnemis pygmaea*.

(DOCX)

